# The temporal and contextual stability of activity levels in hippocampal CA1 cells

**DOI:** 10.1101/2022.01.24.477445

**Authors:** Yuichiro Hayashi, Ko Kobayakawa, Reiko Kobayakawa

## Abstract

Recent long-term optical imaging studies have demonstrated that the activity levels of hippocampal neurons in a familiar environment change on a daily to weekly basis. However, it is unclear whether there is any time-invariant property in the cells’ neural representations. In this study, using miniature fluorescence microscopy, we measured the neural activity of the mouse hippocampus in four different environments every 3 days. Although the activity level of hippocampal neurons fluctuated greatly in each environment across days, we found a significant correlation between the activity levels for different days, and the correlation was higher for averaged activity levels across multiple environments. When the number of environments used for averaging was increased, a higher activity correlation was observed. Furthermore, the number of environments in which a cell showed activity was preserved. Cells that showed place cell activity in many environments had greater spatial information content, and thus carried a higher amount of information about the current position. In contrast, cells that were active only in a small number of environments provided sparse representation for the environment. These results suggest that each cell has not only an inherent activity level but also play a characteristic role in the coding of space.

**Significance Statement:** Recent studies have revealed that place cell activity in the hippocampal CA1 cells exhibit instability on a daily to weekly scale. However, it is unclear whether there is any invariant property in the activity of the cells. In this study, we found that, although the activity level of CA1 neurons fluctuated greatly in one environment, the mean activity level across multiple environments was more stable. Furthermore, the number of environments in which a cell showed activity was preserved over time. These results suggest that even though the spatial code changes dynamically, each cell has an inherent activity level and plays a characteristic role in spatial coding.

## Introduction

The hippocampus is a brain region that plays a crucial role in episodic memory formation. To support this, the activity of hippocampal neurons encode place, environment, time, and various task-relevant information (O’Keefe and Dostrovsky, 1971; Muller and Kubie, 1987; Wood et al., 2000; Ferbinteanu and Shapiro, 2003; Pastalkova et al., 2008; MacDonald et al., 2011). In particular, place cell activity, i.e., the activity of hippocampal cells in restricted portions in an environment, has been extensively studied (O’Keefe and Nadel, 1978; Moser et al., 2015). The activity pattern of place cells can change or disappear when animals move to a different environment (O’Keefe and Conway, 1978; Muller and Kubie, 1987; Alme et al., 2014; Lu et al., 2015). This phenomenon is known as remapping, and it seems to provide a mechanism through which the hippocampus can represent different environments.

Recent studies have shown that the place fields and activity levels of hippocampal neurons for familiar environments are not constant; rather, they fluctuate on a daily to weekly basis (Mankin et al., 2012; Ziv et al., 2013; Rubin et al., 2015; Cai et al., 2016; Hainmueller and Bartos, 2018; Kinsky et al., 2018a; Gonzalez et al., 2019; Hayashi, 2019; Wirtshafter and Disterhoft, 2022). Such fluctuations are known as “representational drift” and have also been found in other brain regions, including the parietal cortex, visual cortex, and olfactory cortex (Driscoll et al., 2017; Deitch et al., 2021; Schoonover et al., 2021). Several studies, however, have provided evidence that each cell has a preconfigured activity level and physiological role; for example, the firing rates of the neurons in the hippocampus and entorhinal cortex are preserved across different brain states and environments (Hirase et al., 2001; Mizuseki and Buzsáki, 2013; Liberti et al., 2022). Recording in a very large environment (>40 m long) revealed that the location of the place fields of the hippocampal neurons fluctuated on a daily to weekly timescale, but that the number of place fields was preserved over months (Lee et al., 2020). Another previous study demonstrated that contextual fear memory engram cells, defined as neurons that excite and express c-fos at memory encoding and whose reactivation results in memory recall, in the CA1 region showed higher environmental specificity and lower spatial information content than non-engram cells in the same area (Tanaka et al., 2018).

In an attempt to reconcile these seemingly incompatible observations on the long-term dynamics of hippocampal activity, we measured neural activity in the mouse hippocampal CA1 area in four different environments over several days. If each cell has a preconfigured activity level and changes its tuning continually, the response to a small set of stimuli (i.e., a small environment) should fluctuate day by day. Conversely, the averaged response to a large variety of stimuli (i.e., a very large environment or multiple environments) will be more stable than that for a single environment. We found that the activity level of hippocampal neurons fluctuated greatly in one environment, and that there was a significant correlation between the activity levels for different days. When the number of environments used for averaging increased, a higher activity correlation was observed. We also found that the number of environments in which the cells showed activity tended to be conserved over days.

## Materials and Methods

### Animal subjects

All experiments complied with the applicable guidelines and regulations. The animal care and use procedure followed protocols that have been approved by the Animal Research Committee of Kansai Medical University. All experiments complied with the ARRIVE guidelines (https://arriveguidelines.org/arrive-guidelines/experimental-animals).

Nine adult male C57BL/6N mice (Japan SLC, Inc., Shizuoka, Japan) aged 12–18 weeks were used in the experiments. The mice were housed under a standard 12-h light/dark cycle and were given access to food and water *ad libitum*.

### Viral constructs

We obtained an AAVrh10.hSyn.GCaMP6f.WPRE.SV40 virus from Vector Core at the University of Pennsylvania at a titer of ∼1.3 × 10^14^ GC/mL. The virus was diluted to ∼5 × 10^12^ GC/mL with phosphate buffered saline before use.

### Miniature head-mount microscopy

The miniature microscope design was based on the UCLA miniscope v.3 (miniscope.org). Excitation light was emitted by a blue LED (LXML-PB01-0030; Lumileds, Aachen, Germany). The light passed through an excitation filter (ET470/40x; Chroma Technology, Bellows Falls, VT) and was reflected by a dichroic mirror (T495lpxr; Chroma Technology) onto the tissue through an electrowetting lens (EWTL) (Varioptic A-25H0; Corning, Corning, NY) and an objective lens (tandem configuration of achromatic doublets; #84-126 and #49-271, Edmund Optics, Burlington, NJ). The fluorescent emissions collected by the lenses were passed through the dichroic mirror and an emission filter (ET525/50m; Chroma Technology). The fluorescence image was focused by a tube lens (#49-277; Edmund Optics) and captured by a CMOS image sensor (MT9V032; ON Semiconductor, Phoenix, AZ). The microscope body was created using a 3D printer (From2, resin type FGPBLK03; Formlabs, Somerville, MA). The design files are available at https://github.com/yuichirohayashi/Zscope. The driving voltage for the EWTL (square wave, 2 kHz) was generated using a data acquisition board (PCIe-6259, National Instruments, Dallas, TX) controlled by custom software written in LABVIEW 7.1 (National Instruments, Dallas, TX) and amplified with a linear amplifier (As-904-150B, NF Corporation, Yokohama, Japan).

### Surgery

Animals were anesthetized using isoflurane. The skull was exposed, and a small hole (<0.5 mm) was made over the right hemisphere (1.5 mm lateral to the midline, 2.3 mm posterior to the bregma). Then, 200 nL of GCaMP6f virus was injected into the CA1 region (1.2 mm ventral from the brain surface). One week after the viral injection, the animals were anesthetized using isoflurane, and a 2.8-mm diameter craniotomy was performed. The dura was removed, and the underlying cortex was aspirated. A stainless-steel cannula (2.76 mm outer diameter, 2.40 mm inner diameter, 1.5 mm height) covered by a glass coverslip (0.12-mm thick) was inserted over the dorsal CA1. Dental cement (Shofu, Kyoto, Japan) was used to glue the cannula to the skull.

Four weeks after the cannula implantation, GCaMP-expressing neurons were imaged using the head-mount microscope. The aluminum baseplate of the microscope was cemented in a position at which the neurons were visible.

### Recording arenas

Calcium activity was recorded in four arenas of different shapes and colors: a tall green box (37 × 25 × 56 cm), a short gray box (37 × 25 × 35 cm), a lime round basket (39 × 26 × 25 cm), and a white square basket (36 × 25 × 25 cm). One side of each arena was painted a different color or decorated with plastic tape (Figure 1A).

**Figure 1.**
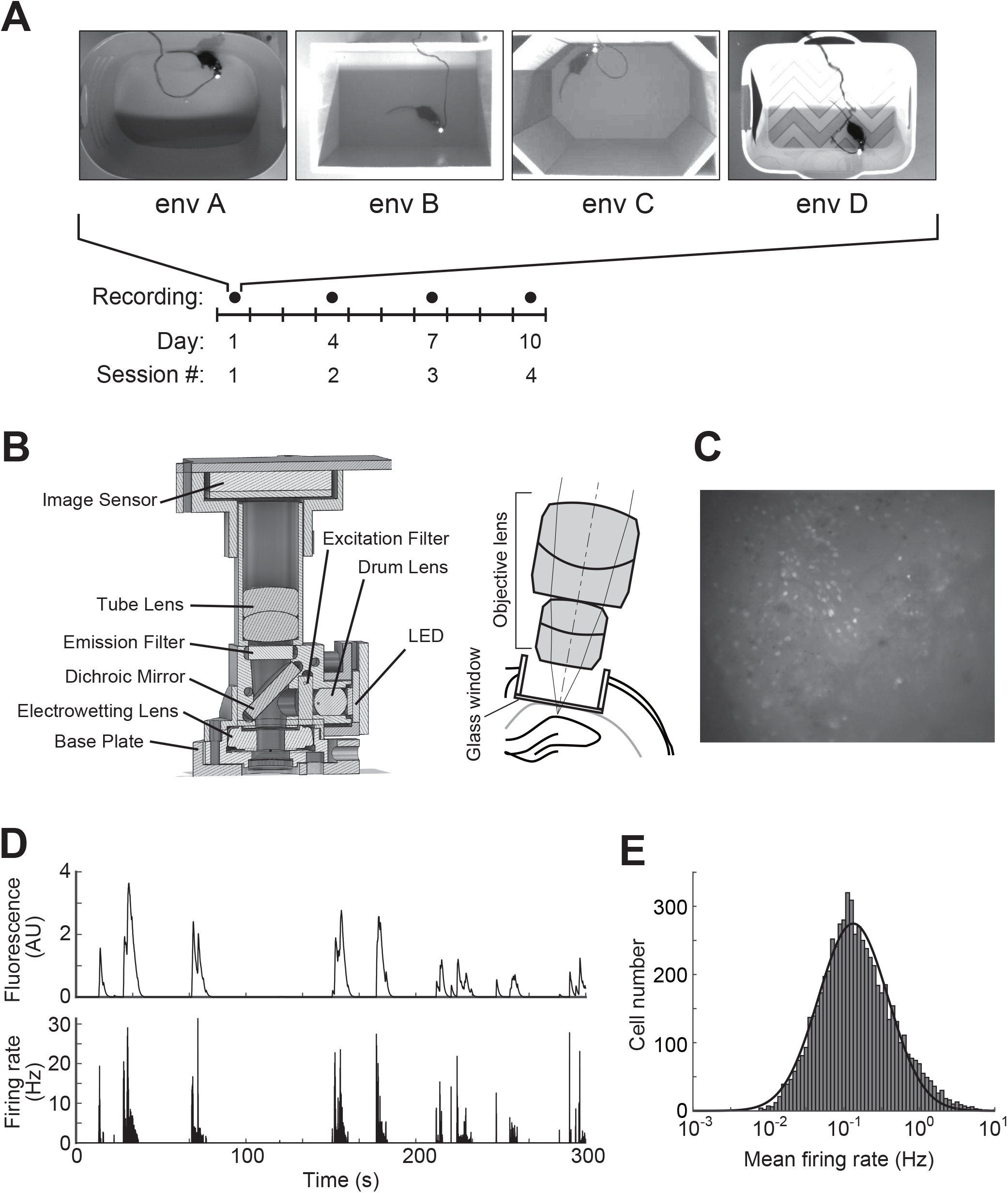
Experimental overview. (**A**) Mice explored four testing arenas per day at 3-day intervals. (**B**) Left: Schematic showing a miniature microscope. Right: Schematic representation showing the objective lens and chronic window implant above the CA1 region. (**C**) Maximum intensity projection of the fluorescence image stream. (**D**) Upper: An example calcium trace of a cell. Lower: Spike train estimated with the OASIS software. (**E**) Firing rate distribution of CA1 cells (n = 6,425 cells). Histogram: data, solid line: lognormal fit.

### Behavioral training

One week after the baseplate surgery, the mice were food-restricted and maintained at above 85% of their initial weight. Mice were trained to forage for food pellets (14 mg, #F05684, Bio-Serv, Flemington, NJ) scattered on the floor of the arenas.

### Recording sessions

After at least 5 days of behavioral training, mice underwent four 5-min recording sessions per day in the four different arenas in a random sequential order. The arenas were cleaned with 70% ethanol prior to each recording session to eliminate any olfactory cues. During the sessions, food pellets were thrown into the arena approximately every 1 min. The excitation light intensity was approximately 0.1–0.5 mW/mm^2^, and fluorescence images were captured at 30 Hz. The focus was adjusted to match the previous image. Animal behavior was recorded at 30 Hz with an overhead camera (DMK22BUC03, The Imaging Source, Bremen, Germany).

### Image processing

ImageJ 1.52 (National Institutes of Health, MA) and MATLAB 2019a (Mathworks, Natick, MA) software was used for all analyses. The calcium imaging data recorded on the same day were concatenated and processed using MiniscopeAnalysis (https://github.com/etterguillaume/MiniscopeAnalysis). First, the images were downsampled spatially by factors of two and motion-corrected using NoRMCorre (Pnevmatikakis and Giovannucci, 2017). The downsampled, motion-corrected videos were then processed using CNMF-E (Zhou et al., 2018) to extract fluorescence traces from individual neurons. The fluorescence traces were finally processed using OASIS software (Pnevmatikakis et al., 2016) for denoizing and deconvolving to estimate the spiking activity. Behavioral image data were binarized such that black pixels represented the animal body. The position of the animal at each frame was determined as the centroid of the black region.

### Registration of cells across sessions

For day-to-day longitudinal cell registration, the spatial footprints of the cells extracted with CNMF-E from each day’s fluorescence image stream were processed with the Cellreg routine (Sheintuch et al., 2017) using the default parameters (maximum distance <12 µm, spatial correlation >0.74, and P_same_ threshold = 0.5). In brief, the sets of spatial footprints of cells for each session were aligned with rigid-body translation. The distribution of spatial footprint similarities between pairs of neighboring cells from different sessions was modeled to obtain an estimation for the probability of them being the same cell (P_same_). Registration quality was assessed using the average P_same_ value for registered cells in an animal and the register score for registered cells (Sheintuch et al., 2017). The register score was calculated as the total number of reliably identified cell pairs out of the total number of cell pairs across all sessions using the following equation:

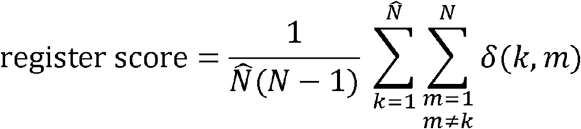

where N represents the total number of sessions, ∼N represents the number of active sessions for a given cell register. The *δ*(*k,m*) value was 1 if a cell-pair in the cluster was reliable and 0 if the cell-pair was unreliable.

### Place fields

To analyze the place fields, calcium events that occurred when the animal’s velocity was less than 1 cm/s were filtered out to eliminate nonspecific activity at rest. The place field maps were constructed by finding the spike number at each spatial bin (2 × 2 cm) divided by the dwell time in the bin. The maps were smoothed using a Gaussian function with a standard deviation of 3 cm.

For each place field of the cells, the mutual information between calcium transients and the mouse’s location was calculated (Skaggs et al., 1993). To assess the significance of spatial selectivity, Monte Carlo *P* values were calculated. A total of 1,000 distinct shuffles of the calcium transient times were performed, and the mutual information for each shuffle was calculated. The *P* value was defined as the fraction of shuffles that exceeded the mutual information of the cell. Cells with a *p* value of <0.05 were considered place cells.

### Rate difference

The rate difference was determined by calculating the unsigned rate difference between the mean calcium transient rates in the two recording sessions and then dividing the difference by the sum of the two rates.

### Population vector correlation

To compute the population vector (PV) correlation between two sessions, the place fields of all simultaneously recorded place cells were stacked into a 3D matrix (place field map × cell identity). At each spatial bin, the firing rate vector of all cells represented the PV for that spatial bin. To compare two recording sessions, the Pearson’s correlation coefficient was calculated between each pair of PVs at the corresponding locations, and the correlation coefficients of all spatial bins were averaged (Leutgeb et al., 2005).

### Experimental design and statistical analysis

Calcium activity was recorded from nine mice. Recordings from animals with low activity (average arena coverage <80%) were discarded. Statistical analysis was performed using GraphPad Prism 5 (GraphPad Software, La Jolla, CA), MATLAB 2019a, and R version 4.1.1. Data were first tested for normality using the Shapiro–Wilk test and for homogeneity of variance with Bartlett’s test or the *F*-test. Normally distributed data with equal variances were compared with a one-way analysis of variance (ANOVA) followed by Bonferroni’s post hoc comparisons. If data were not normally distributed, nonparametric tests (Mann–Whitney U test, Kruskal–Wallis test, or Friedman test) were used. Statistical significance was set at *p* < 0.05 for all statistical analyses.

## Results

### Experimental outline

To investigate the place cell activity for multiple spatial environments in the hippocampus, neural activity was recorded successively in four arenas with different shapes and colors (Figure 1A). Before commencing the recording sessions, mice (n = 6) were trained to find food in these arenas for at least 5 days. Each mouse then underwent four 5-min recording sessions per day at 3-day intervals (Figure 1A).

Neural activity in the dorsal CA1 region of the hippocampus was recorded using a combination of a genetically encoded calcium indicator, GCaMP6f, and a miniature microscope (Figure 1B, C). We extracted the calcium traces of putative neurons from the fluorescence image stream using CNMF-E (Zhou et al., 2018). The firing rate of each neuron was estimated from its calcium trace using OASIS software (Pnevmatikakis et al., 2016) (Figure 1D). The firing rate distribution of the recorded cells showed lognormal-like distribution (Figure 1E), consistent with the findings of a previous electrophysiological study (Mizuseki and Buzsáki, 2013). With the spatial footprints of the cells extracted with CNMF-E from each day’s fluorescence image stream, we used Cellreg (Sheintuch et al., 2017) to track cells across days.

### Activity of CA1 cells showed a variety of environmental specificity

A total of 6,425 cells were recorded from six mice (602–1,580 cells per animal) in four different environments. Following a previous study, we observed various cells ranging from those that showed no activity in any environment to those that showed activity in all four environments in one day (Figure 2A, B) (Lu et al., 2015). We termed the number of environments where a cell exhibited activity above the threshold as the “global activity level index” (gA index). We also classified cells according to the number of environments where a cell showed statistically significant place cell activity. This value was termed the “global place cell activity level index” (gPC index). As shown in Figure 2C, CA1 cells displayed various gPC activity levels. However, a fraction of cells that showed significant place cell activity did not change between environments or over days, indicating that these mice were highly familiar with all the environments (Figure 2D, Q_(3,69)_ = 1.639, *p* = 0.65, n = 24 recordings per day in six animals. Q_(3,69)_ = 5.812, *p* = 0.12, n = 24 recordings for each environment in six animals, Friedman test).

**Figure 2.**
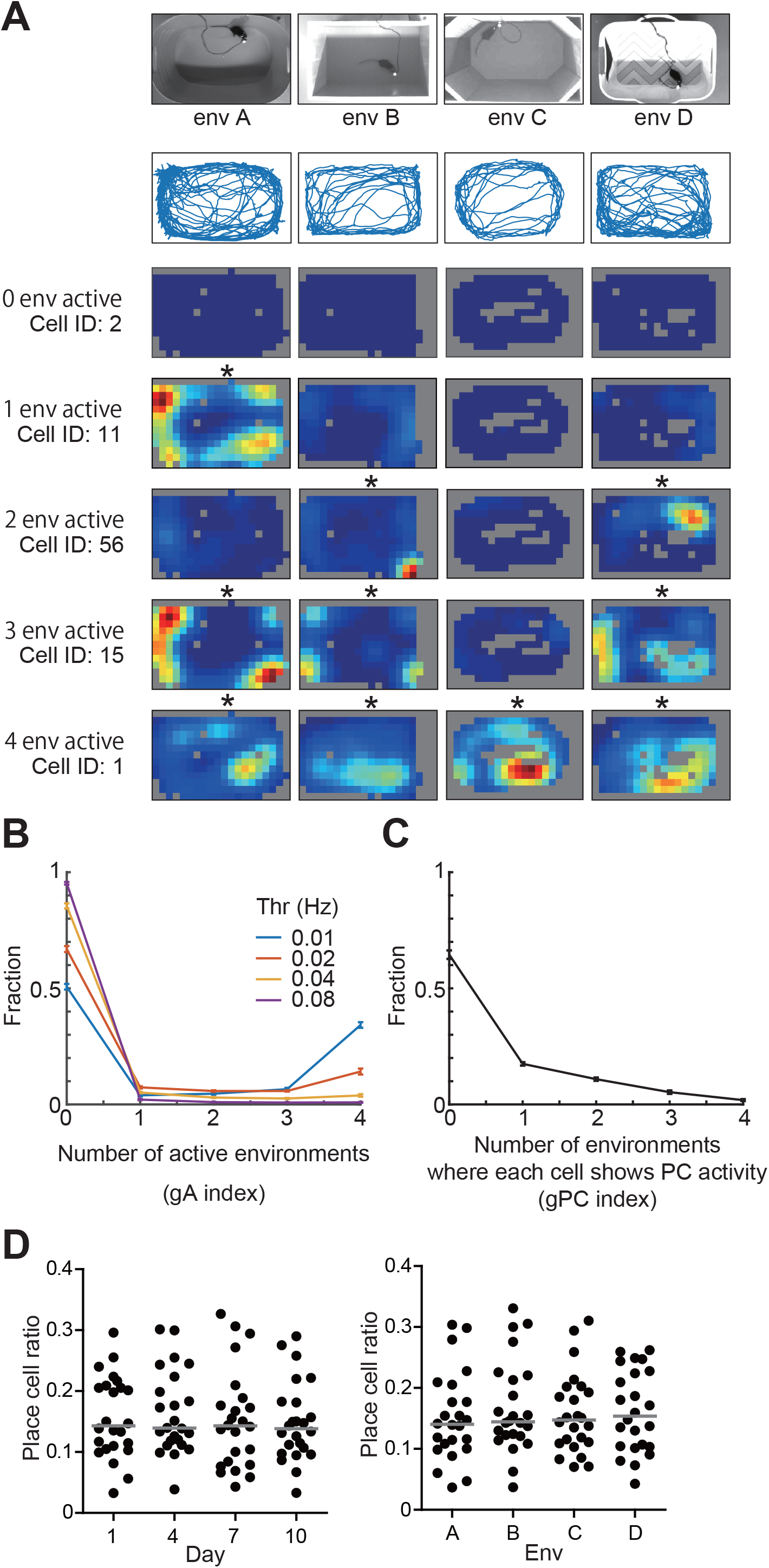
Hippocampal activity recorded in the four environments. Place cell activity in four different environments. (**A**) Top row: Top view showing the four environments. Second row: Typical mouse trajectories. Remaining rows: Place field maps for five representative cells. Red indicates the maximum firing rate, blue indicates silence. Regions not visited by the mice are shown in gray. Asterisks represent activity above the threshold (mean firing rate > 0.01 Hz). (**B**) Distribution of the gA index with a variable threshold for activity. Data are presented as the mean ± SEM (n = 24 sessions involving 6 mice). (**C**) Distribution of the gPC index. Data are presented as the mean ± SEM (n = 24 sessions involving 6 mice). (**D**) Place cell ratio on each day (left) and in each environment (right). Each dot shows an individual recording in a single environment. Solid lines indicate the mean value.

### Long-term tracking of neural activity for each cell

For longitudinal analysis of the same cells, the cells need to be reliably registered across days. Therefore, we checked the register score and the P_same_ value to ensure the reliability of cell registration (Sheintuch et al., 2017). The average registration score was 0.86 ± 0.020 (mean ± SEM, n = 6 animals, Figure S1A, left), and the average P_same_ value was 0.83 ± 0.052 (mean ± SEM, n = 6 animals, Figure S1A, right). These values were comparable to those in the previous study in the hippocampal CA1 area (Sheintuch et al., 2017).

Next, we evaluated the reliability of cell registration at the functional level. For this purpose, the PV correlation (Leutgeb et al., 2005) of the spatial representation for the four environments between sessions was calculated. Consistent with previous studies (Rubin et al., 2015; Kinsky et al., 2018b), the PV correlation between sessions in the same environment was significantly higher than between sessions in different environments for at least a 6-day session interval (Figure S1B, *F*_(5,174)_ = 5.302, *p* < 0.0001, n = 24 session pairs for same environment, n = 36 session pairs for different environments, one-way ANOVA followed by Bonferroni’s post hoc test). The results also corroborated that the same neurons were reliably tracked across multiple sessions.

Not only across-day but also within-day registration quality affects the analysis results. For wide-field imaging data, signals from neighboring cells overlap due to light scattering and lack of optical sectioning capability (Wilt et al., 2009). Furthermore, although CNMF-E has been shown to effectively demix signals from overlapping cells (Zhou et al., 2018), a possible concern remains that the activities of multiple spatially overlapping cells may be merged into the activity of a single high-gA cell. Therefore, to test this possibility, we examined the relationship between a cell’s gA index and its spatial footprint size of the region of interest extracted using CNMF-E. If multiple spatially overlapping cells are merged into a single cell, its spatial footprint is hypothesized to be larger than that for a genuinely single cell. As shown in Figure S1C, there was a slight negative correlation between the gA index and the spatial footprint size. This result suggested that the demix error in the cellular signal extraction was not more frequent for high-gA cells than for low-gA cells.

### Activity levels averaged across multiple environments were more stable than those for a single environment

Next, we analyzed the long-term dynamics of neural activity for each cell in the four different environments. A significant correlation between the activity levels in an environment for adjacent sessions (3 days apart) was observed (Figure 3A, “1 env”, R = 0.66, n = 77,100 observations of cells). Notably, the correlation for mean activity level across multiple environments increased gradually according to the increased number of environments to be averaged. For calculating the firing rate for multiple environments, to match the data length, we used 5-min data constructed by concatenating data fragments randomly chosen from recordings in multiple environments (Figure 3A, “2 env”, R = 0.69, n = 115,650 observations of cells, “3 env”, R = 0.70, n = 77,100 observations of cells, “4 env”, R = 0.71, n = 19,275 observations of cells; Figure 3B, *F*-_(3,15)_ = 24.70, *p* < 0.0001, n = 6 animals, one-way repeated measures ANOVA followed by Tukey’s post hoc test).

**Figure 3.**
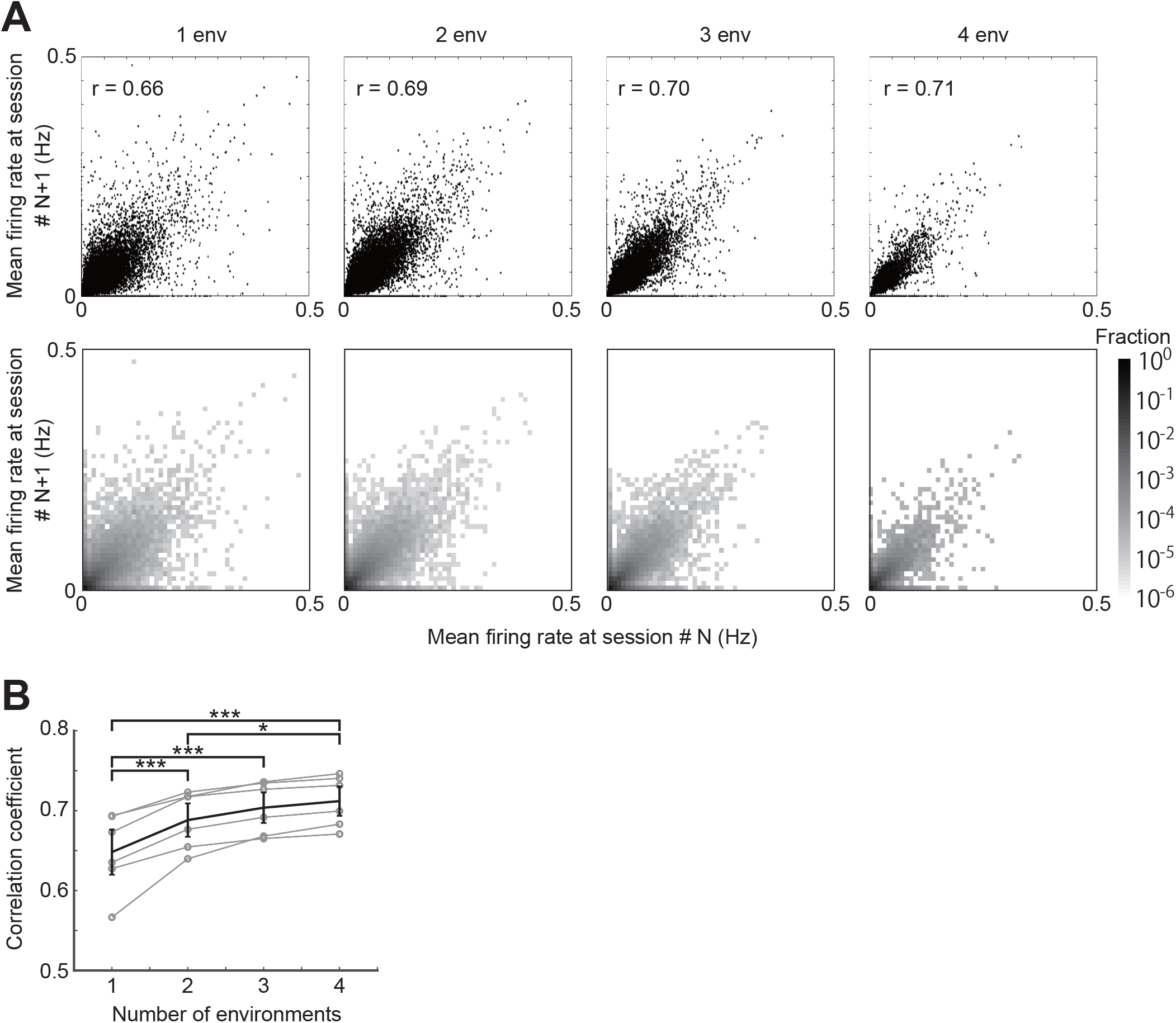
Firing rate correlation across sessions. (**A**) Change in firing rate between two adjacent sessions. Each dot in the upper panels shows one cell’s firing rate across two adjacent sessions. Heat maps in the lower panels show the fraction of cells. Left to right: columns show the firing rates in a single (same) environment, average firing rates in 5-min data randomly gathered from 2, 3, and 4 environments. Pearson’s correlation coefficient (R) is displayed in each panel. (**B**) The correlation coefficients for individual animals. The black line shows mean ± SEM. Each gray circle shows an individual animal (n = 6 animals). Asterisks indicate significant differences (\**p* < 0.05, \*\*\**p* < 0.0001).

### The gA and gPC indices were preserved over time

We then tested whether the cells’ response properties to environments, such as the gA and gPC indices, were also preserved. A significant correlation was observed for the gA index between adjacent sessions (Figure 4A, n = 90 session pairs, R = 0.78, *p* < 0.0001). Furthermore, although we used a threshold value of 0.01 Hz, a significant correlation was still found with threshold levels ranging from 0.01 to 0.08 Hz (Figure S2). The gPC index between adjacent sessions was also correlated (Figure 4B, n = 90 session pairs, R = 0.72, *p* < 0.0001). These results indicated that not only the mean firing rate but also the gA and gPC indices (gA/gPC indices) of each cell were preserved over time.

**Figure 4.**
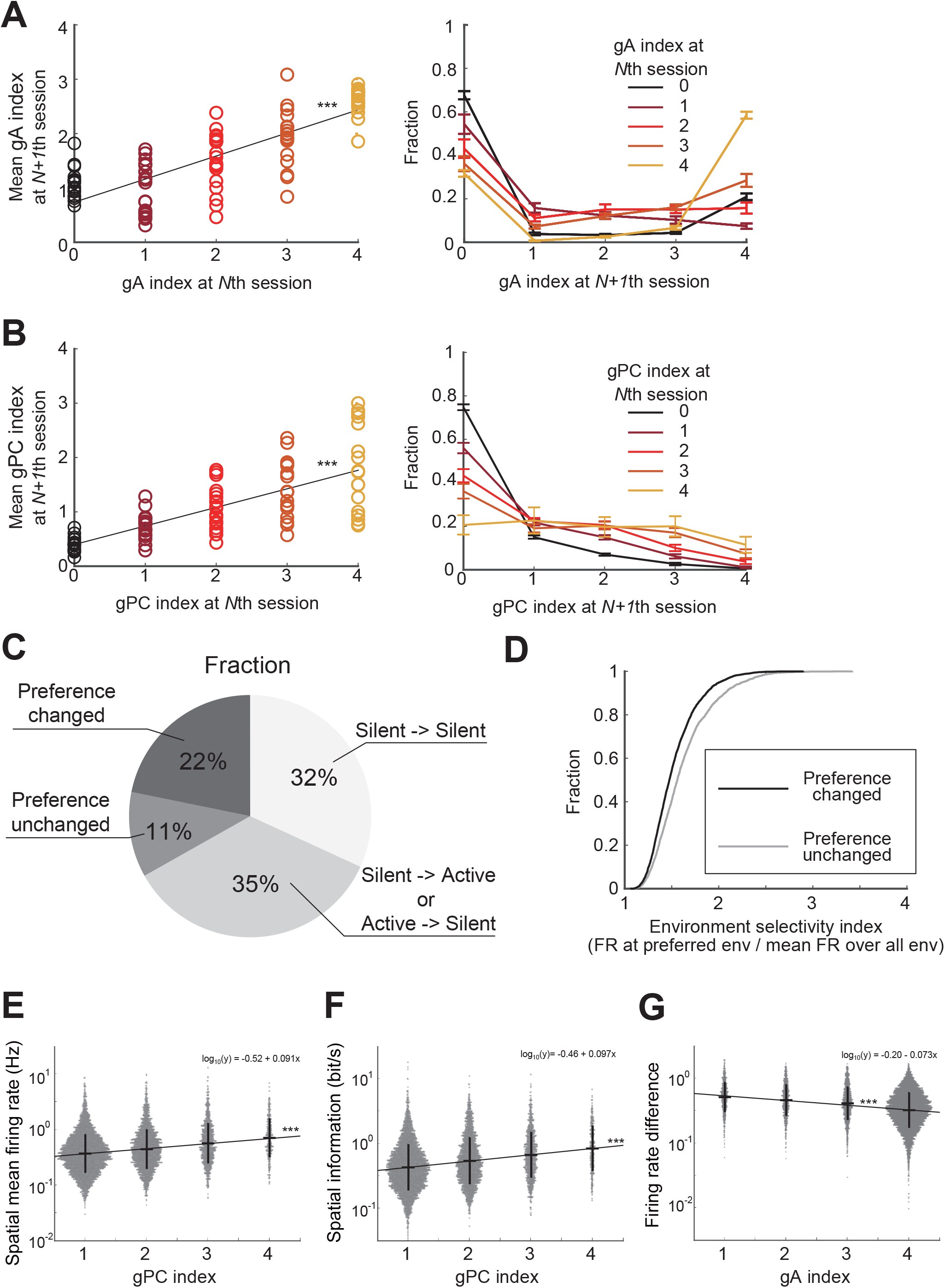
Session-to-session dynamics of the gA/gPC indices. **(A)** Changes in cell’s gA index between two adjacent sessions. Left: Open circles represent each value. The solid line shows a linear regression fit to the data. Right: Distribution of gA index at *N+1*st session for cells showing each gA index at *N*th session. Data are presented as mean ± SEM (n = 18 session pairs from six animals). **(B)** Changes in cell’s gPC index between two adjacent sessions. Left: Open circles represent each value. The solid line shows a linear regression fit to the data. Right: Distribution of gPC index at *N+1*st session for cells showing each gA index at *N*th session. Data are presented as mean ± SEM (n = 18 session pairs from six animals). Solid line shows a linear regression fit to the data. Asterisks indicate significant correlations (\*\*\**p* < 0.0001). (**C**) Fraction of “preference changed” cells and “preference unchanged” cells between sessions out of 19,275 observations of the cells. Others showed no activity in either or both day (“silent->silent” or “silent->active or active->silent”, respectively). (**D**) Cumulative distribution of environment preference index for “preference changed” cells and “preference unchanged” cells. (**E**) The gPC index versus the mean firing rates. The solid line shows a linear regression fit to the data (log-transformed). (**F**) The gPC index versus the spatial information rates. The solid line shows a linear regression fit to the data (log-transformed). (**G**) gA index versus the firing rate difference for the environment in which a cell showed the highest firing rate and firing rate averaged over the other three environments. The solid line shows a linear regression fit to the data (log-transformed). Dots represent each value. Horizontal lines show the mean values, and the whiskers show the standard deviation. Asterisks indicate significant differences (\*\*\**p* < 0.0001).

Although the gA/gPC indices were preserved over time, a significant fraction of the cells changed their environmental preferences (Figure 4C, D). Additionally, on the basis that each cell had preserved activity levels and gA/gPC indices, we determined whether each cell also had an inherent role in spatial coding. We investigated the relationship between the gA/gPC indices and the cells’ firing properties. Both the mean firing rates and spatial information rates were positively correlated with the cell’s gPC index (Figure 4E, mean firing rate: R = 0.22, F_(3, 8514)_ = 424, *p* = 5.3 × 10^−92^, n = 8,516 observations of neurons over all recording sessions; 6B, spatial information: R = 0.23, F_(3, 8514)_ = 467, *p* = 6.9 × 10^−101^, n = 8,516 observations of place cells over all recording sessions, *F*-test). These results indicated that high gPC index cells convey much information about the current position. In contrast, cells that were active in a small number of environments constituted a sparse code for environmental identity. To confirm this, we calculated the firing rate differences between the environments where a cell showed the highest firing rate and the firing rate averaged over the other three environments. Because cells did not necessarily possess position information when encoding the environment, we classified the cells using the gA index. The firing rate difference was negatively correlated with the gA index (Figure 4G, R = 0.24, F_(3,12310)_ = 761, *p* = 1.6 × 10^−162^, n = 12,312 observations of cells over all recording sessions, *F*-test). This result indicated that cells with a small gA index show sparse representation for environment.

## Discussion

In this study, using a head-mount miniature fluorescence microscope, we measured the neural activity of hippocampal CA1 cells over time in four different environments (Figure 1A). We found a significant correlation for each cell’s activity level for a single environment between sessions, and the correlation for mean activity level across environments increased gradually according to the increased number of environments to be averaged (Figure 3A, B). This result reconciles previous conflicting observations on the long-term stability of the neuronal activity in the CA1: the place field location and activity levels of CA1 cells are not constant but fluctuate on a daily to weekly basis (Ziv et al., 2013; Rubin et al., 2015; Cai et al., 2016; Kinsky et al., 2018a, 2020; Gonzalez et al., 2019; Hayashi, 2019). In contradiction to these reports, a recent study reported that the activity levels of CA1 cells in a very large (∼40 m long) environment were stable over time, though the location of their place field fluctuated (Lee et al., 2020). If each cell has a preconfigured activity level and changes its tuning continually, the response to a small set of stimuli will fluctuate day by day. Conversely, the averaged response to a large variety of stimuli will be stable. Animals receive a larger variety of sensory stimuli from a large environment or multiple environments than from a single environment. This is probably the reason why averaged activity across multiple environments is stable over days (Figure 3A, B).

We also showed that the gA and gPC indices tended to be preserved over time (Figure 4A, B). This suggests that each CA1 cell possesses not only a preconfigured activity level but also an inherent role in spatial coding. Cells with a high gPC index convey a higher amount of spatial information per unit of time; that is, these cells contribute more to coding their location in an environment than those with a low gPC index (Figure 4F). In contrast, cells with a low-gA index sparsely code the environmental identity. Cells with diverse gA/gPC indices in the CA1 cooperatively code external space with multiresolution. The sparse code for environment allows simple mechanisms for forming associations with specific environments (Olshausen and Field, 2004). In the CA1, contextual fear memory engram cells, defined as neurons that excite and express c-fos at memory encoding and whose reactivation results in memory recall, show high environmental specificity and low spatial information content (Tanaka et al., 2018). The activity profile of the memory engram cells resembles that of low-gA cells in the present study (Figure 4F). Therefore, it would be interesting to examine the relationship between a cell’s c-fos expression during spatial memory encoding and its gA/gPC indices.

Although each cell’s activity level and gA/gPC indices were preserved over time, we also showed that the environmental tuning of its activity was unstable (Figure 4C, D). However, at the population level, CA1 cells showed distinct representations for different environments for at least 6 days (Figure S1B). These results suggest that environmental information was retained as population activity in CA1 cells.

We employed calcium imaging for long-term tracking of individual cell activity, which cannot be accomplished reliably using electrophysiological techniques. For this technique, however, the neocortex above the hippocampus needs to be removed for optical access to the CA1 cells (Figure 1B). Although the basic activity profiles of the CA1 cells were consistent with those reported in previous electrophysiological studies (Figure 1E, 2A), an unknown effect of this surgery on their neural activity cannot be excluded. Validation of our results will be obtained using a less invasive optical method (Kondo et al., 2017).

To drive GCaMP expression, we used human synapsin I gene promoter, which promotes transgene expression in both excitatory and inhibitory cells. Recent studies have reported that a subset of inhibitory cells in the CA1 show spatial and environmental modulation in activity (Geiller et al., 2020; Schuette et al., 2022). Therefore, it will be of interest and value to study the relationship between gA/gPC indices and subtypes of neurons.

The studies of long-term hippocampal activity measurement discussed here were performed in mice. However, recent application of calcium imaging on species other than mice have revealed different activity profiles of CA1 cells. For example, highly stable activity and spatial tuning were detected in bat CA1 cells, and a high percentage of place cells were identified in rat CA1 cells (Liberti et al., 2022; Wirtshafter and Disterhoft, 2022). The temporal dynamics of gA/gPC indices may also be different in these species.

## Abbreviations

AAV: adeno-associated virus
GC: genome copy
ANOVA: analysis of variance
EWTL: electrowetting lens
PV: population vector

## Acknowledgments

This work is supported by JSPS KAKENHI (Grant No. 17K19436, 20K07715) to Y.H., JSPS KAKENHI 20H04849 to R.K, and AMED-CREST 21GM1510003 to K.K. The authors would like to thank Enago (www.enago.jp) for the English language review.

