## Supplementaly material for "The temporal and contextual stability of activity levels in hippocampal CA1 cells"

This PDF file includes:

Fig S1-2

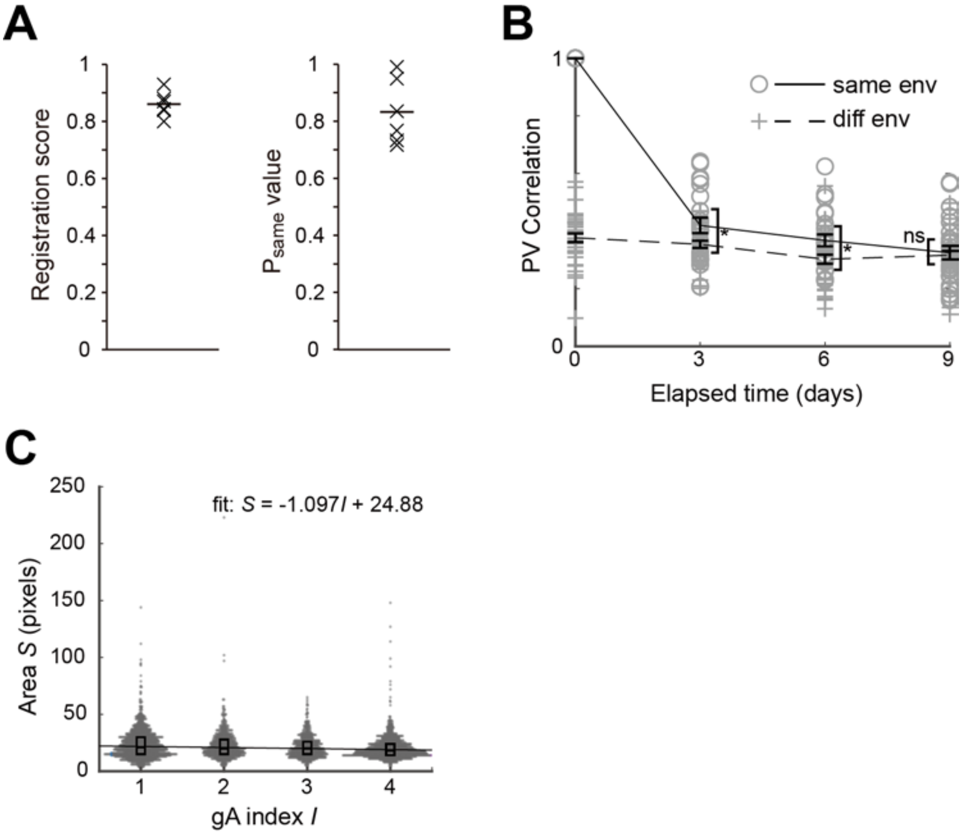


**Figure S1. Cell registration across sessions**

(**A**) Distribution of registration scores (left) and average $P_{same}$ values (right) for all animals (n = 6 animals). Each crossbar represents an individual animal. Solid lines indicate the mean value. (**B**) Population vector (PV) correlation for the same (solid line) and different (dashed line) environments across time. Data are presented as mean ± SEM. Open circles represent each value (n = 24 session pairs) for the same environment, cross markers represent each value (n = 36 session pairs) for different environments. Statistical significance was tested using a two-tailed *t*-test with Bonferroni's correction. Asterisks indicate significant differences (*$p < 0.05$, ns: not significant). (**C**) Each cell's gA index versus the spatial footprint sizes of their spatial components extracted by CNMF-E. The spatial footprint size is the number of pixels whose value is greater than the half-maximal of the cell' s spatial components. Dots represent each value. Box plots show the median and quartiles. The solid line shows a linear regression fit to the data. R = 0.112, $p < 0.0001$.

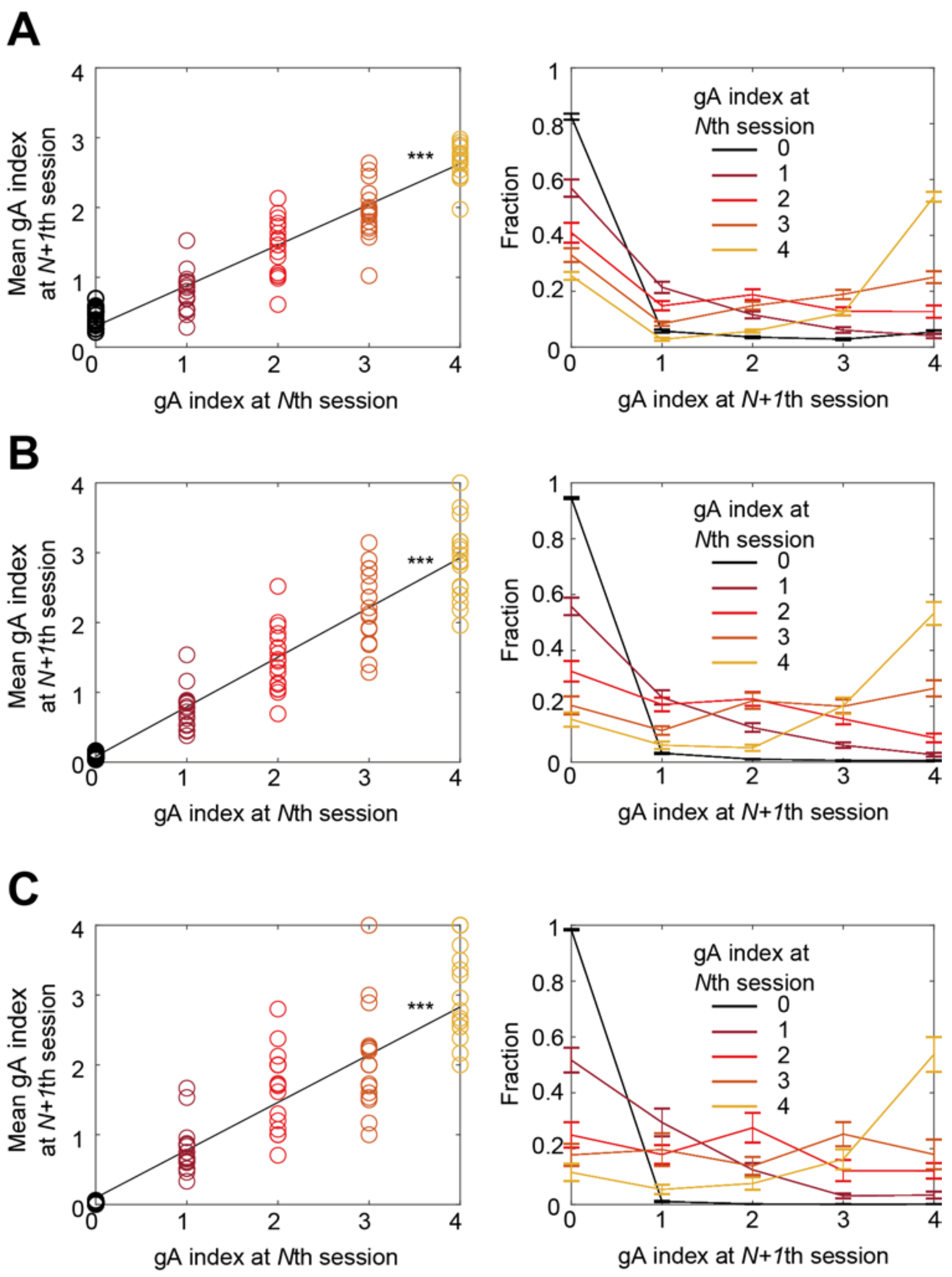


**Figure S2. Session-to-session dynamics of gA index with variable threshold**

(**A**) Changes in gA index between two adjacent sessions. Activity threshold = 0.02 Hz. n = 90 session pairs, r = 0.934, $p < 0.0001$. (**B**) Changes in gA index between two adjacent sessions. Activity threshold = 0.04 Hz. n = 90 session pairs, r = 0.929, $p < 0.0001$. (**C**) Changes in gA index between two adjacent sessions. Activity threshold = 0.08 Hz. n = 90 session pairs, r = 0.772, $p < 0.0001$

.
